# Phenotypic expansion in *DDX3X* – a common cause of intellectual disability in females

**DOI:** 10.1101/283598

**Authors:** Xia Wang, Jill A. Rosenfeld, Carlos A. Bacino, Fernando Scaglia, LaDonna Immken, Jill M. Harris, Scott E. Hickey, Theresa M. Mosher, Anne Slavotinek, Jing Zhang, Joke Beuten, Magalie S. Leduc, Weimin He, Francesco Vetrini, Magdalena A. Walkiewicz, Weimin Bi, Rui Xiao, Yunru Shao, Alper Gezdirici, Yunyun Jiang, Adam W. Hansen, Davut Pehlivan, Juliette Piard, Donna M. Muzny, Neil Hanchard, John W. Belmont, Lionel van Maldergem, Richard A. Gibbs, Mohammad K. Eldomery, Zeynep C. Akdemir, Tamar Harel, Jennifer E. Posey, Adekunle M. Adesina, Shan Chen, members of the UDN, Brendan Lee, James R. Lupski, Christine M. Eng, Fan Xia, Yaping Yang, Brett H. Graham, Paolo Moretti

## Abstract

*De novo* variants in *DDX3X* account for 1-3% of unexplained intellectual disability (ID), one of the most common causes of ID, in females. Forty-seven patients (44 females, 3 males) have been described. We identified 29 additional individuals carrying 27 unique *DDX3X* variants in the setting of complex clinical presentations including developmental delay or ID. In addition to previously reported manifestations, rare or novel phenotypes were identified including respiratory problems, congenital heart disease, skeletal muscle mitochondrial DNA depletion, and late-onset neurologic decline. Our findings expand the spectrum of DNA variants and phenotypes associated with *DDX3X* disorders.

## Introduction

Intellectual disability (ID) affects 1-3% of the population, with higher prevalence in males versus females^1^. In males pathogenic variants in over 100 genes on the X chromosome account for 5-10% of ID^2,3^. Relatively less is known about the number of X-linked genes causing ID in females^4^, but whole exome sequencing (WES) is finding *de novo* variants in X-linked ID genes in females of all ages^5–7^. However, limited information is available regarding such cases. Some of the genes causing ID in females are known to cause disease in males, including *PHF6*^8^ and *USP9X*, with the latter causing congenital malformations not observed in affected males^9^. Further evidence for gender-specific variant pathogenicity comes from *DDX3X* located on Xp11.4, with pathogenic *de novo* variants causing syndromic ID in 39 females; in the same study, three males inherited *DDX3X* variants from apparently unaffected mothers^10^. Differences in predicted variant severity or X-chromosome inactivation studies from blood DNA did not explain the gender-specific disease expression. Five additional females with *DDX3X* variants have been described in the literature^11–13^. These reports led us to hypothesize that females with *de novo* variation in *DDX3X* may show additional clinical phenotypes. We report 29 individuals with *DDX3X*-related disorders, and provide comprehensive clinical presentations for ten, including the oldest reported individual. These data expand the number of *DDX3X* pathogenic variants and their associated phenotypic spectrum.

## Methods

Variants in *DDX3X* were identified by WES, performed according to previously described methods^5,6,12^, either on a clinical basis at Baylor Genetics (Females 1-24, Males 1-2) or on a research basis by the Baylor Hopkins Center for Mendelian Genomics (BHCMG, Females 25-27). De-identified reporting of demographic and molecular data for all clinically-referred cases was approved by the Institutional Review Board at Baylor College of Medicine (BCM). Additional, informed consent for publication of clinical details was obtained for a subset of clinically-referred cases and all research-based cases according to IRB-approved protocols: at Baylor College of Medicine (Female 8, 14, 17, 23, 24), through the Undiagnosed Diseases Network (UDN) protocol (Female 13), and through the BHCMG (Females 7, 19, 25, 27). Female 7 and 19 were previously reported in a study of research-based reanalysis of clinical WES data^12^. *DDX3X* variants were annotated using transcript NM_001193416. Variant pathogenicity was determined based on the ACMG guidelines^14^ and the internal guidelines developed at Baylor Genetics.

## Results

Among 4839 (2152 females, 2687 males) patients referred to the Baylor Genetics laboratory for clinical WES with developmental delay (DD) and/or ID, 26 (24 females, 2 males) were found to carry pathogenic or likely pathogenic variants in *DDX3X*. Through collaboration with the BHCMG, an additional 3 unrelated female cases were identified. The ages at molecular diagnosis ranged from 1 to 47 years (Table S1). Twenty-seven unique variants were identified (24 novel and 3 reported previously), including 11 missense, 6 frameshift, 3 splice site, 4 nonsense, and 3 in-frame deletion/duplication changes (Table 1 and Figure 1A and B). In 27 individuals with available parents (26 female and 1 male), the *DDX3X* variants were confirmed as *de novo*, supporting the variant pathogenicity. Of the 27 *de novo* mutations, 7 occurred at CpG dinucleotides. Two *de novo* variants, c.573_575del (p.I191del) and c.1805G>A (p.R602Q) are mosaic in the proband, with allele fractions of 21% and 14%, respectively (Table 1).

**Table 1.**
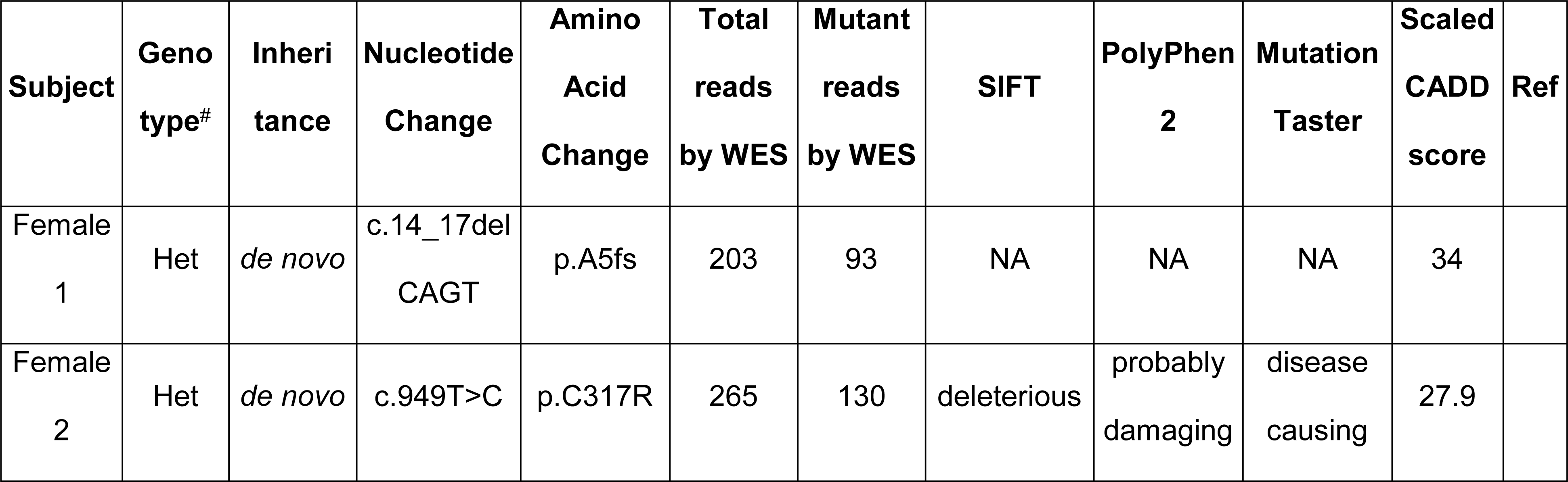
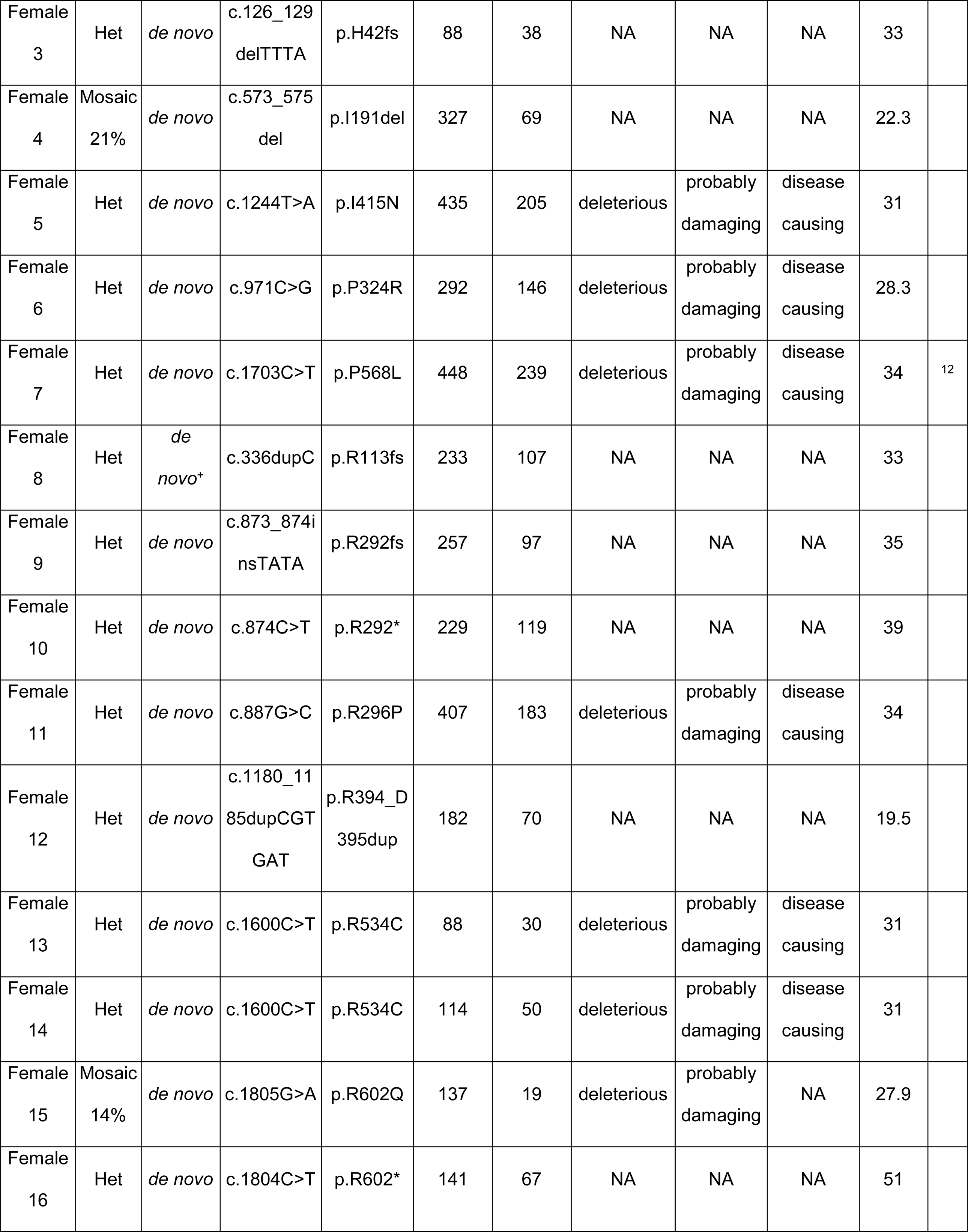
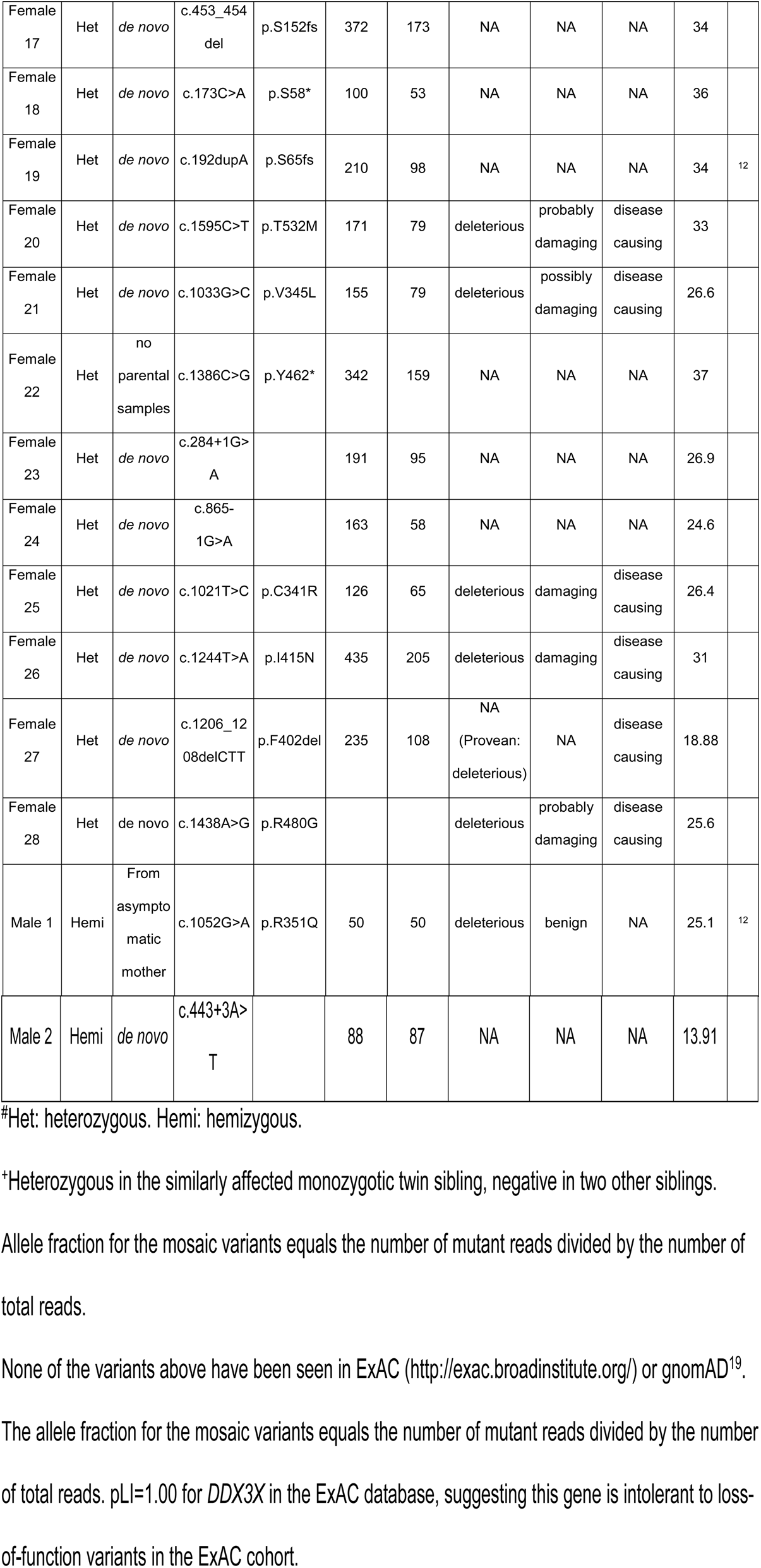
Subjects with pathogenic or likely pathogenic variants in *DDX3X*.

| Subject | Genotype<sup>#</sup> | Inheritance | Nucleotide Change | Amino Acid Change | Total reads by WES | Mutant reads by WES | SIFT | PolyPhen 2 | Mutation Taster | Scaled CADD score | Ref |
| --- | --- | --- | --- | --- | --- | --- | --- | --- | --- | --- | --- |
| Female 1 | Het | *de novo* | c.14_17del<br>CAGT | p.A5fs | 203 | 93 | NA | NA | NA | 34 |  |
| Female 2 | Het | *de novo* | c.949T>C | p.C317R | 265 | 130 | deleterious | probably damaging | disease causing | 27.9 |  |
| Female<br>3 | Het | *de novo* | c.126_129<br>delTTTA | p.H42fs | 88 | 38 | NA | NA | NA | 33 |  |
| Female<br>4 | Mosaic<br>21% | *de novo* | c.573_575<br>del | p.I191del | 327 | 69 | NA | NA | NA | 22.3 |  |
| Female<br>5 | Het | *de novo* | c.1244T>A | p.I415N | 435 | 205 | deleterious | probably<br>damaging | disease<br>causing | 31 |  |
| Female<br>6 | Het | *de novo* | c.971C>G | p.P324R | 292 | 146 | deleterious | probably<br>damaging | disease<br>causing | 28.3 |  |
| Female<br>7 | Het | *de novo* | c.1703C>T | p.P568L | 448 | 239 | deleterious | probably<br>damaging | disease<br>causing | 34 | 12 |
| Female<br>8 | Het | *de*<br>*novo*<sup>+</sup> | c.336dupC | p.R113fs | 233 | 107 | NA | NA | NA | 33 |  |
| Female<br>9 | Het | *de novo* | c.873_874i<br>nsTATA | p.R292fs | 257 | 97 | NA | NA | NA | 35 |  |
| Female<br>10 | Het | *de novo* | c.874C>T | p.R292* | 229 | 119 | NA | NA | NA | 39 |  |
| Female<br>11 | Het | *de novo* | c.887G>C | p.R296P | 407 | 183 | deleterious | probably<br>damaging | disease<br>causing | 34 |  |
| Female<br>12 | Het | *de novo* | c.1180_11<br>85dupCGT<br>GAT | p.R394_D<br>395dup | 182 | 70 | NA | NA | NA | 19.5 |  |
| Female<br>13 | Het | *de novo* | c.1600C>T | p.R534C | 88 | 30 | deleterious | probably<br>damaging | disease<br>causing | 31 |  |
| Female<br>14 | Het | *de novo* | c.1600C>T | p.R534C | 114 | 50 | deleterious | probably<br>damaging | disease<br>causing | 31 |  |
| Female<br>15 | Mosaic<br>14% | *de novo* | c.1805G>A | p.R602Q | 137 | 19 | deleterious | probably<br>damaging | NA | 27.9 |  |
| Female<br>16 | Het | *de novo* | c.1804C>T | p.R602* | 141 | 67 | NA | NA | NA | 51 |  |
| Female<br>17 | Het | *de novo* | c.453_454<br>del | p.S152fs | 372 | 173 | NA | NA | NA | 34 |  |
| Female<br>18 | Het | *de novo* | c.173C>A | p.S58* | 100 | 53 | NA | NA | NA | 36 |  |
| Female<br>19 | Het | *de novo* | c.192dupA | p.S65fs | 210 | 98 | NA | NA | NA | 34 | 12 |
| Female<br>20 | Het | *de novo* | c.1595C>T | p.T532M | 171 | 79 | deleterious | probably<br>damaging | disease<br>causing | 33 |  |
| Female<br>21 | Het | *de novo* | c.1033G>C | p.V345L | 155 | 79 | deleterious | possibly<br>damaging | disease<br>causing | 26.6 |  |
| Female<br>22 | Het | no<br>parental<br>samples | c.1386C>G | p.Y462* | 342 | 159 | NA | NA | NA | 37 |  |
| Female<br>23 | Het | *de novo* | c.284+1G><br>A |  | 191 | 95 | NA | NA | NA | 26.9 |  |
| Female<br>24 | Het | *de novo* | c.865-1G>A |  | 163 | 58 | NA | NA | NA | 24.6 |  |
| Female<br>25 | Het | *de novo* | c.1021T>C | p.C341R | 126 | 65 | deleterious | damaging | disease<br>causing | 26.4 |  |
| Female<br>26 | Het | *de novo* | c.1244T>A | p.I415N | 435 | 205 | deleterious | damaging | disease<br>causing | 31 |  |
| Female<br>27 | Het | *de novo* | c.1206_12<br>08delCTT | p.F402del | 235 | 108 | NA<br>(Provean:<br>deleterious) | NA | disease<br>causing | 18.88 |  |
| Female<br>28 | Het | *de novo* | c.1438A>G | p.R480G |  |  | deleterious | probably<br>damaging | disease<br>causing | 25.6 |  |
| Male 1 | Hemi | From<br>asympto<br>matic<br>mother | c.1052G>A | p.R351Q | 50 | 50 | deleterious | benign | NA | 25.1 | 12 |
| Male 2 | Hemi | *de novo* | c.443+3A><br>T |  | 88 | 87 | NA | NA | NA | 13.91 |  |

#Het: heterozygous. Hemi: hemizygous.

**Figure 1.**
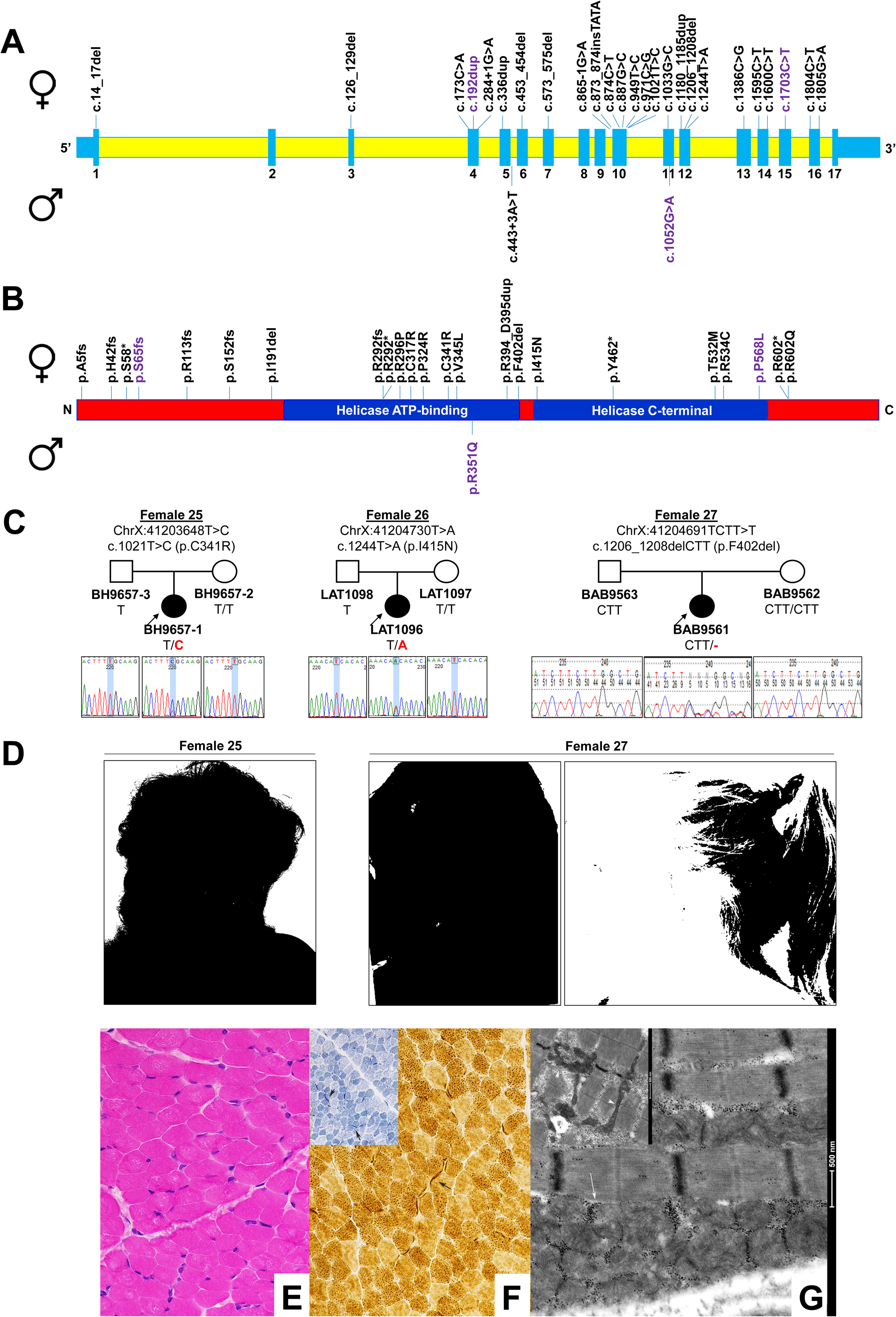
Location of *DDX3X* variants identified in this study, Female individuals (25, 26, 27) ascertained through the BHCMG, and muscle biopsy results in Female 17 showing abnormal mitochondrial morphology. (A) Schematic view of the *DDX3X* exon-intron structure based on NM_001193416. Blue boxes represent exons and yellow fields represent introns. Exon number is listed below each exon. cDNA change is listed for each variant. (B) Schematic view of the DDX3X protein structure based on Snijders Blok et al. Amino acid change is listed for each variant. Variant color: black, first reported in this study; purple, previously reported. (C) Pedigree and Sanger tracings demonstrate *de novo* inheritance in three unrelated female probands. (D) Female 25 demonstrated synophrys, a broad nasal root with upturned nostrils, a long philtrum, and thin upper lip. Female 27 demonstrated cupped ears, a long philtrum, and a thin upper lip. (E) Skeletal muscle cross-section showing mild variation in fiber size (H&E; magnification x400). (F) Skeletal muscle cross-section showing few fibers with mild subsarcolemmal increase in oxidative activity [cytochrome oxidase (long arrow) and NADH tetrazolium reductase (insetarrow heads; magnifications x400)]. (G) Electron microscopic images showing mild subsarcolemmal mitochondrial proliferation (long arrow) with inset in the right upper corner showing pleomorphic abnormally elongated and irregularly shaped mitochondria (arrow heads).

The most frequent clinical presentations in the 27 females include DD and/or ID (27/27), hypotonia (19/27), dysmorphic features (19/27), structural brain abnormalities (17/19 who had brain MRI), movement disorders (16/27), visual impairments (8/27), and microcephaly (7/27) (Table 2). Some individuals had respiratory problems (5/24): apnea, tachypnea, and chronic respiratory failure. Congenital heart disease (5/7 who had echo) included atrial/ventricular septal defect, double orifice mitral valve with small patent ductus arteriosus, mild concentric left ventricular hypertrophy and bicuspid aortic valve. In comparison to published data, autism spectrum disorder and other behavior problems and skin abnormalities are underrepresented in our cohort (Table 2). For ten subjects, we obtained additional informed consent and provide detailed clinical descriptions (Table S1), as well as clinical images for two subjects (Figure 1C and D).

**Table 2.** Comparison of clinical presentations in this study and in the published cohort.

| **Clinical features** | **Number of subjects**<br>**in this study** | **Percentage in this**<br>**study** | **Percentage in the**<br>**published cohort** |
| --- | --- | --- | --- |
| DD and/or ID | 27/27 | 100% | 100% |
| Hypotonia | 19/27 | 70% | 76% |
| Dysmorphic features | 19/27 | 70% | NA |
| Structural brain<br>abnormalities | 17/19 | 89% | 81% |
| Movement disorders | 16/27 | 59% | 45% |
| Visual impairments | 8/27 | 30% | 34% |
| Microcephaly | 7/27 | 26% | 32% |
| Autism spectrum<br>disorders and other<br>behavior problems | 6/27 | 22% | 53% |
| Respiratory problems | 5/24 | 21% | NA |
| Congenital heart disease | 5/7 | 71% | NA |
| Skin abnormalities | 4/27 | 15% | 37% |

NA: not specified or reported in the published study<sup>10</sup>.
In comparison to published data, autism spectrum disorder and other behavior problems and skin abnormalities are underrepresented in our cohort: 6/27 vs 20/38 ( $P=6.8 \times 10^{-3}$ ) and 4/27 vs 14/38 ( $P=2.5 \times 10^{-2}$ ), respectively.

In two subjects, skeletal muscle mitochondrial DNA content was reduced. The first subject (Female 17) is a 6 year-old non-dysmorphic girl with a history of neonatal hypotonia, esophageal reflux and global developmental delay. A quadriceps muscle biopsy demonstrated mild fiber type variation and abnormal pleomorphic mitochondria on electron microscopy (Figure 1E-G). Analysis of respiratory chain enzyme activities demonstrated reductions of multiple complexes, with relative sparing of complex II activity and reduction of citrate synthase activity. Sequencing of mitochondrial DNA from the muscle sample did not detect any known or likely pathogenic variants. Mitochondrial DNA content in muscle was 39% of age-matched control muscle. Clinical WES demonstrated a *de novo* heterozygous c.453_454del (p.S152fs) pathogenic variant in *DDX3X*, with no other pathogenic or likely pathogenic variants identified in known disease associated genes. The second subject is a 47-year-old woman (Female 13) with history of global developmental delay, short stature, dysmorphic features, microcephaly and unilateral renal agenesis. She learned to sit at two years of age and walk at eight, and she only learned to say simple words. In her early 40s, she regressed, becoming non-verbal and unable to ambulate or to use her arms. She was found to carry a *de novo* heterozygous c.1600C>T (p.R534C) likely pathogenic variant in *DDX3X* (Table 1). The same variant was also observed in Female 14 in our cohort, and a variant involving the same codon (p.R534H) has been reported in a patient with ID^10^. A quadriceps muscle biopsy demonstrated severe mitochondrial and lipid depletion, and reduction in mitochondrial size similar to Female 17. Mitochondrial DNA content in muscle was 26% of the mean value for age and tissue matched controls. A reduction in all mitochondrial respiratory chain complex activities and citrate synthase activity was also observed.

## Discussion

Normal RNA metabolism requires the function of RNA helicases (RH), and yet, the exact function of most human RH remains unknown. Genetic studies have begun to address the role of altered RH function in human disease^15^. *DDX3X* encodes a DEAD-box RNA helicase important in transcription, splicing, RNA transport, and translation^16,17^. In a diagnostic laboratory referral cohort of 4839 subjects with DD and/or ID, we have identified 26 individuals (24 females, 2 males) with syndromic ID or DD carrying pathogenic or likely pathogenic variants in *DDX3X*; an additional 3 females were identified through research WES at BHCMG. The overall frequency of pathogenic or likely pathogenic *DDX3X* variation in our cohort is 0.54% of the total (26/4839) and 1.12% of females (24/2152), similar to a previous report (0.6% and 1.5% respectively)^10^, confirming *DDX3X* as one of the most common genetic causes of unexplained ID in females. In our laboratory, *DDX3X* ranks third among approximately 450 genes for the occurrence of *de novo* variants, with *ARID1B* first (43 patients) and *ANKRD11* second (29 individuals). In addition to confirming published data, our phenotypic analysis expands the phenotypic spectrum associated with *DDX3X* variants in females. For instance, we found respiratory problems and congenital heart disease in 5/24 and 5/7 of our subjects, phenotypes not previously described in the original description of *DDX3X* related disorders^10^, although observed in a subsequent report of two females^11^. We found no evidence for genotype-phenotype correlations between the mutations we identified and age at onset or phenotypic severity. Previously reported individuals ranged in ages from 1 to 33 years. We report the phenotype of a 47-year-old woman (Female 13) who has manifestations consistent with *DDX3X* disorder and a clinical picture of previously unreported late-onset neurologic decline. The decline is unrelated to intercurrent illness, and her motor function is at least in part responsive to physical therapy. In two individuals (Female 13 and 17), we found skeletal muscle mitochondrial DNA depletion, in a range similar to what is observed in patients with proven mitochondrial DNA depletion syndrome. However, neither case carried pathogenic variants in genes associated with this phenotype. This observation is relevant for at least two reasons. First, if confirmed in additional subjects, it may provide an explanation for the mechanism of some of the observed phenotypes, including the late-onset neurologic decline seen in the oldest of our subjects. Second, *DDX3X* dysfunction may need to be considered in individuals with unexplained mitochondrial DNA depletion. Although the mechanism of mitochondrial DNA depletion in these individuals remains to be elucidated, pathogenic variants in *DDX3X* have been shown to regulate Wnt signaling-mediated ventralization in females^10^. Interestingly, Wnt signaling has also been shown to regulate mitochondrial biogenesis^18^. Molecular studies in cell culture and model organisms will be required to elucidate the role of *DDX3X* variants in regulating mitochondrial function and abundance. Two variants reported in this study, c.1600C>T (p.R534C) and c.1703C>T (p.P568L), and three previously reported, c.641T>C (p.I214T), c.931C>T (p.R311*), and c.1084C>T (p.R362C)^10^, have also been seen somatically in medulloblastoma, malignant melanoma, and esophageal squamous cell carcinoma (http://cancer.sanger.ac.uk/cosmic). Malignancy has not been reported in the 29 patients included in this study. However, pathway analysis for the highest 5% and lowest 5% genes expressed in RNAseq data from dermal fibroblasts obtained in one subject (Female 13) showed enrichment in cell cycle control of chromosomal replication and double-strand break repair pathways (data not shown). Future studies will elucidate whether individuals carrying *DDX3X* variants are at risk for the development of malignancies. In summary, we identified 29 unrelated patients with causal variants in *DDX3X* and expanded the genotypic and phenotypic spectrum of *DDX3X* related disorders. The collective data suggest that *DDX3X* defects are a frequent cause of syndromic ID in females, and the causal variants are likely to be loss-of-function.

## Acknowledgements

We are deeply grateful to the patients and clinicians whose participation made this study possible. Research reported in this manuscript was supported by the NIH Common Fund, through the Office of Strategic Coordination/Office of the NIH Director under Award Number U01HG007709, and the National Human Genome Research Institute (NHGRI)/National Heart Lung and Blood Institute (NHLBI) UM1 HG006542 to the Baylor Hopkins Center for Mendelian Genomics (JRL). The content is solely the responsibility of the authors and does not necessarily represent the official views of the National Institutes of Health. JEP was supported by K08 HG008986 through the National Human Genome Research Institute.

## Author Contributions

XW: acquisition and analysis of data, and drafting a significant portion of the manuscript or figures

JAR: acquisition and analysis of data, and drafting a significant portion of the manuscript or figures

CAB: acquisition and analysis of data

FS: acquisition and analysis of data

LI: acquisition and analysis of data

JMH: acquisition and analysis of data

SEH: acquisition and analysis of data

TMM: acquisition and analysis of data

AS: acquisition and analysis of data

JZ: acquisition and analysis of data

JB: acquisition and analysis of data

MSL: acquisition and analysis of data

WH: acquisition and analysis of data

FV: acquisition and analysis of data

MAW: acquisition and analysis of data

WB: acquisition and analysis of data

RX: acquisition and analysis of data

YS: acquisition and analysis of data

AG: acquisition and analysis of data

YJ: acquisition and analysis of data

AWH: acquisition and analysis of data

DP: acquisition and analysis of data

JP: acquisition and analysis of data

DMM: acquisition and analysis of data

NH: acquisition and analysis of data

JWB: acquisition and analysis of data

LVM: acquisition and analysis of data

RAG: acquisition and analysis of data

MKE: acquisition and analysis of data

ZCA: acquisition and analysis of data

TH: acquisition and analysis of data

JEP: acquisition and analysis of data, and drafting a significant portion of the manuscript or figures

AMA: acquisition and analysis of data

SC: acquisition and analysis of data

BL: conception and design of the study, acquisition and analysis of data, and drafting a significant portion of the manuscript or figures

JRL: conception and design of the study, acquisition and analysis of data, and drafting a significant portion of the manuscript or figures

CME: acquisition and analysis of data

FX: acquisition and analysis of data

YY: acquisition and analysis of data

BHG: conception and design of the study, acquisition and analysis of data, and drafting a significant portion of the manuscript or figures

PM: conception and design of the study, acquisition and analysis of data, and drafting a significant portion of the manuscript or figures

## Figure Legends

**Table S1.** (see excel file) Clinical features and *DDX3X* variants in subjects enrolled in this study. Detailed clinical features are only reported for the subjects in whom we were able to obtain additional informed consent.

